# Empirical Bayes Meets Information Theoretical Network Reconstruction from Single Cell Data

**DOI:** 10.1101/264853

**Authors:** Thalia E. Chan, Ananth V. Pallaseni, Ann C. Babtie, Kirsten R. McEwen, Michael P.H. Stumpf

## Abstract

Gene expression is controlled by networks of transcription factors and regulators, but the structure of these networks is as yet poorly understood and is thus inferred from data. Recent work has shown the efficacy of information theoretical approaches for network reconstruction from single cell transcriptomic data. Such methods use information to estimate dependence between every pair of genes in the dataset, then edges are inferred between top-scoring pairs. Dependence, however, does not indicate significance, and the definition of “top-scoring” is often arbitrary and a *priori* related to expected network size. This makes comparing networks across datasets difficult, because networks of a similar size are not necessarily similarly accurate. We present a method for performing formal hypothesis tests on putative network edges derived from information theory, bringing together empirical Bayes and work on theoretical null distributions for information measures. Thresholding based on empirical Bayes allows us to control network accuracy according to how we intend to use the network. Using single cell data from mouse pluripotent stem cells, we recover known interactions and suggest several new interactions for experimental validation (using a stringent threshold) and discover high-level interactions between sub-networks (using a more relaxed threshold). Furthermore, our method allows for the inclusion of prior information. We use *in-silico* data to show that even relatively poor quality prior information can increase the accuracy of a network, and demonstrate that the accuracy of networks inferred from single cell data can sometimes be improved by priors from population-level ChIP-Seq and qPCR data.

## Introduction

In all organisms gene expression appears to be carefully controlled. Such control is achieved by networks of transcription factors and a whole host of regulators (Trapnell et al., 2014; Gouti et al., 2015; Göttgens, 2015). These act in concert to determine when — and in higher organisms, also where — genes are turned on or off. The processes of gene expression, together with those of degradation, activation, and, where relevant, transport, determine the abundance and activity of proteins inside cells.

The nature and structure of these networks is only incompletely understood (Young et al., 2014; Huang and Zi, 2014). Even for the most studied model organisms we only have outlines and rough representations of these networks. For higher organisms, we are becoming ever more aware how inadequate the simple network models of the past are: regulation is a dynamical, and furthermore dynamically controlled process involving a large number of different components, that together orchestrate gene expression.

Technological advances notwithstanding, full elucidation of these extensive and complex networks using principled and comprehensive experimental analysis will remain impossible for the foreseeable future. Instead we require a combination of experiments and statistical inference to tackle this problem. Statistical inference in this context is primarily concerned with the *detection* of potential candidate relationships between, for example, gene expression measurements (here typically quantitative measurements of mRNA) of different genes. These may indicate the existence of either regulatory interactions (between a regulatory controller, e.g. a transcription factor, and a target), or co-regulatory interactions (where two targets are controlled by the same transcription factor).

Biological network inference has been an active field for as long as high-throughput experimental technologies have been available (Penfold and Wild, 2011; Penfold et al., 2015; Bonneau et al., 2006; Villaverde et al., 2013; Vinciotti et al., 2016), and the recent advent of single cell transcriptomic data has given rise to a resurgence (Babtie et al., 2017). Despite this, the different techniques (and, indeed, combinations of approaches (Marbach et al., 2012; Villaverde et al., 2015)) have typically only been applied to the current data at hand, without paying attention to any available prior knowledge such as previously inferred network models. Thus the effort has largely been on making best use of the available data, but scant attention has been given to available background information; some notable exceptions exist (Mukherjee and Speed, 2008; Olsen et al., 2014; Studham et al., 2014).

Information theoretical (IT) approaches have gained prominence as tools for network reconstruction complementing an array of other approaches from statistics and control engineering (Krishnaswamy et al., 2014; Zhao et al., 2016; Villaverde et al., 2016; Chan et al., 2017). They make minimal assumptions; have desirable Statistical properties (especially in situations where different scenarios should be *a priori* weighted equally probable) (Cover and Thomas, 2012; Mc Mahon et al., 2014; Kinney and Atwal, 2014); and they seem to be particularly promising for many applications in molecular and cellular biology, where noise and potential non-linearities are expected to be pervasive (Mc Mahon et al., 2015; Uda et al., 2013). Current single cell transcriptomic data challenge many of the existing network reconstruction approaches due to technical and biological heterogeneity (as reviewed elsewhere (Babtie et al., 2017)). Recent applications, e.g. in the context of gene regulatory network reconstruction, are beginning to suggest that the information theoretical framework is ideal for dealing with single cell transcriptomic data (Chan et al., 2017). But it has been notoriously difficult to standardize the interpretation of these information theoretical measures, since they depend on the size and treatment of the dataset and hence vary widely between studies (Altay and Emmert-Streib, 2010; Chan et al., 2017).

Here we attempt to address this problem: we show how it is possible to employ an *empirical Bayes* (EB) approach in the statistical interpretation of information theoretical measures in the context of network inference. This has two advantages: first, we can apply formal hypothesis testing/model selection criteria in order to determine significant relationships; and second, we can introduce a *post hoc* adjustment to the EB prior in order to incorporate any available prior information (fig. 1).

**Figure 1:**
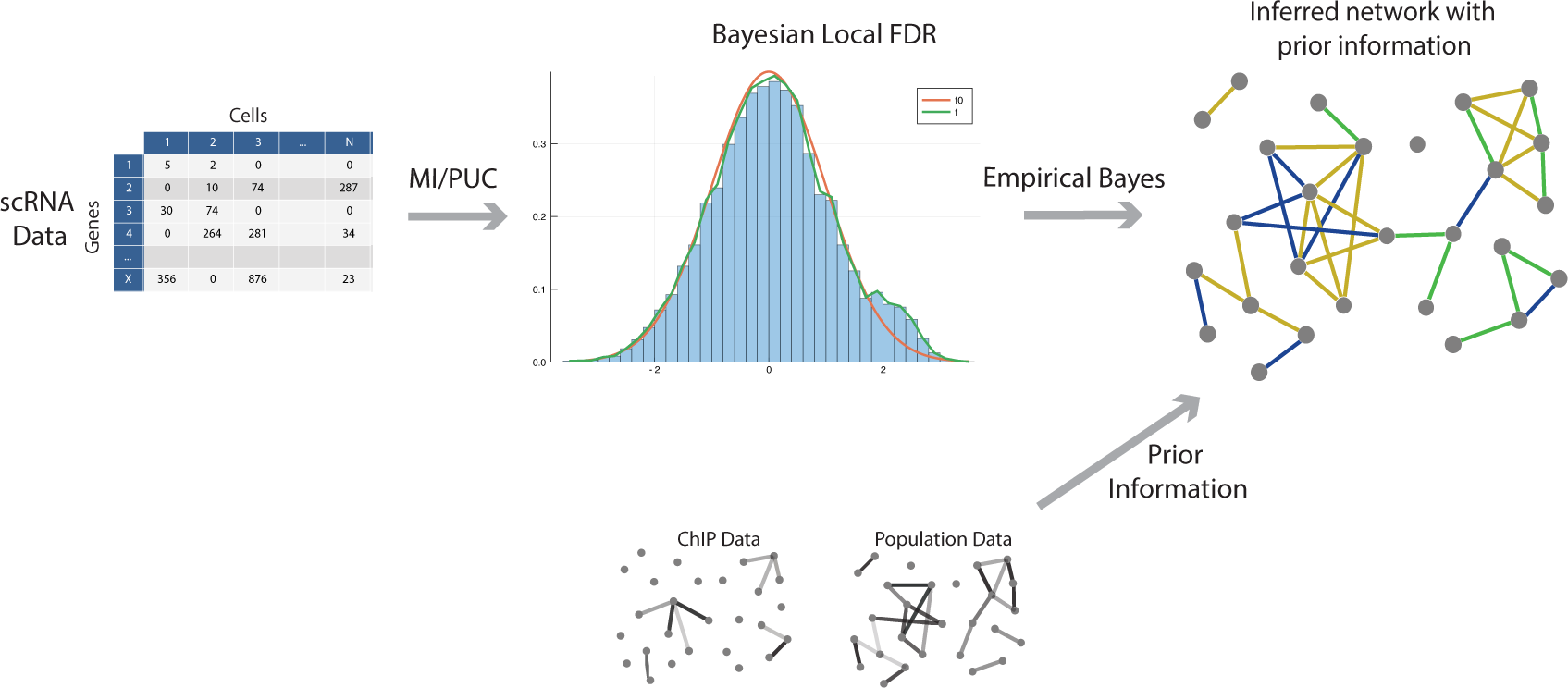
Using data, e.g. from single cell qPCR or RNA-seq, and any available prior information, our ap-proach uses the Empirical Bayes local False Discovery Rate (FDR) to infer edges (i.e. statical dependencies that could signal potential functional relationships) among sets of genes.

Below we introduce information theoretical measures used in network inference, before outlining the EB framework, and our testing and evaluation procedure. The advantages of the EB+IT approach become clear in applications to simulated (where we know the true network) and real data: we find that we can control the accuracy of our networks according to the use case. Supplementing experimental data with available background information can improve the performance of network inference compared to the case where only the data in hand are considered, and we will provide some quantitative guidelines as to how reliable prior information has to be in order to make a positive difference. Interestingly, we find that single cell data are so informative - and higher-order information theoretic measures sufficiently powerful - that the best results are sometimes gained using single cell data alone.

## Information Measures for Network Reconstruction

The *mutual information* (Kraskov et al., 2004; Steuer et al., 2002) captures the decrease in the uncertainty about random variable (RV), *X,* if the state of RV *Y* is known (or *vice versa* as *I(X; Y)= I(Y;X))*,

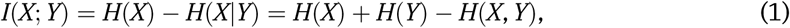

Where *H(X)* and *H(X/Y)* denote the *entropy* of random variable *X*, and the *conditional entropy* of *X* given the state of *Y*, respectively. *I(X; Y)* is the generic measure to detect statistical dependencies between pairs of random variables (e.g. expression levels of genes).

Going beyond pairwise measures of information has proved difficult, but recently the *partial information decomposition* (PID), and related measures have shown considerable promise (Chan et al., 2017; Williams and Beer, 2010). These measures dissect the statistical dependence between a target RV, e.g. *Z,* and a set, *S*, of explanatory variables, e.g. *S = {X, Y},* which can help with filtering out indirect and spurious interactions.

PID is defined as (Williams and Beer, 2010)

*I(Z;X, Y) =* Synergy(*Z;X, Y*) + Unique_Y_*(Z;X)* + Unique_X_*(Z; Y)* + Redundancy*(Z;X, Y),* where *I(Z;X) =* Unique_Y_*(Z;X) +* Redundancy(Z;*X,* Y); with the specific information, as defined by Williams and Beer (Williams and Beer, 2010),

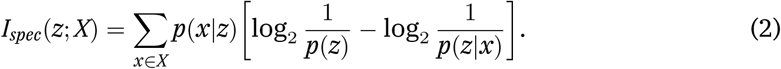

We can write for the different terms in (),

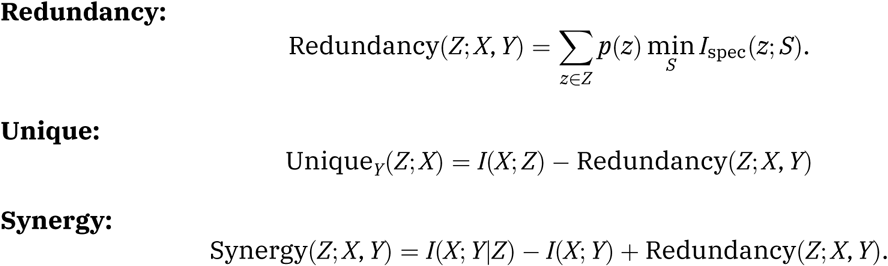

The *proportional unique contribution* (PUC) between two nodes *X* and *Z* is the sum of the ratio of unique information to mutual information, calculated over every other node *Y* in a network (where *S* is the complete set of nodes) (Chan et al., 2017),

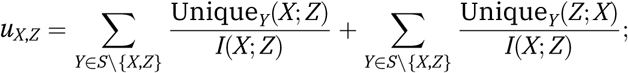

and it forms a promising and powerful way of assessing the relative weight in favor of an interaction between RVs *X* and Z.

MI, PUC, and similar quantities give continuous measures for the statistical dependencies between pairs of nodes in a network. Their interpretation is often made difficult by the fact that these measures are not normalized; nor are they easily linked to statistical significance of edges. Instead heuristics or arbitrary thresholds are used to score edges. A further shortcoming is that this framework does not cater for the inclusion of any *apriori* knowledge (including indirect evidence for the existence) of edges: all that is taken into account is the available data, which although increasingly this may seem plentiful, potentially ignores much of what is already known. Here we use the EB framework to address these problems.

## An Empirical Bayes Framework for Information Theoretical Network Reconstruction

With the increased availability of high-throughput data, it is not uncommon to test hundreds or thousands of hypotheses at once, be it in gene expression analysis or GWAS studies. Hypothesis testing relies on the formulation of a meaningful null distribution, but for many complicated problems these may be hard to come by. Even when we can come up with plausible null hypotheses and distributions, the distribution of the observed test statistics can deviate from the theoretical null, even when it is expected that the null hypothesis is true in the vast majority (e.g. over 90%) of cases. Such deviations can be explained (for example, unobserved covariates may lead to overdispersion (Efron, 2004)), but if ignored may lead to incorrect interpretations of significance.

EB provides a framework for large-scale simultaneous hypothesis testing, in which an empirical null distribution is derived from the observed test statistics, under the assumption that non-null cases are rare. Network inference can be viewed as such a problem: for every pair of genes in the dataset, we have a null hypothesis that there is no interaction (i.e. that vectors of their expression across the cells are statistically independent). Thus, for a dataset of *n* genes, we have 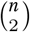 hypotheses, and from the knowledge that we do have about gene regulatory networks (Marbach et al., 2012), we expect the null hypothesis to hold for the vast majority of pairs.

We follow Efron’s approach (Efron et al., 2003; Efron, 2007) and determine the local false discovery rate (FDR), fdr(x), where *x* here denotes the measures of interest (i.e. either the MI or PUC). Values of *x* are drawn from a two component mixture model,

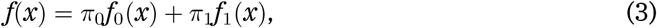

Where *f_0_(x)* and *f_1_(x)* denote the probability densities correspondingto the null and nonnull models, respectively (Efron, 2004). In the present context, to be concrete, all values corresponding to non-interacting/dependent pairs of nodes are assumed to be drawn from a probability model with associated density, *f*_0_.

We seek to determine the local FDR,

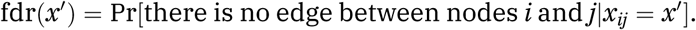

This probability is given, for *N* observations as *π_0_f_0_(x')/f(x'),* which leaves us with the problems of estimating the null, *f_0_(x),* and empirical, *f(x),* densities. Even for medium-sized networks the size of the available IT estimates is sufficiently large for us to estimate these densities from the data (see the following section for details). These are, of course, not statistically independent, but the expectations of the estimates depend only weakly on such correlations in the data; the sampling variance, by contrast, can vary considerably due to the presence of correlations (Efron, 2008).

This yields the estimate and the p-value

This yields the estimate

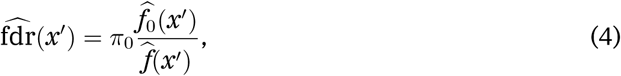

and the p-value

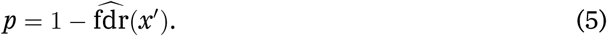

Note that *p* indicates the fraction of tests with MI (or PUC) *x'* expected not to be false discoveries.

## Estimation of Empirical Bayes Distributions

In large-scale hypothesis testing the observed null distribution often diverges from the theoretical null; hence it is often appropriate to use a null of the correct theoretical form, with empirically-derived parameters (Efron, 2004). Even determining the appropriate theoretical null model for information measures is problematic, however.

It has been shown that the MI between independent random variables is gamma distributed (Goebel et al., 2005), and our simulations support this (fig. 2). The theoretical null of PUC has not previously been characterized, but following arguments e.g. in (Hutter, 2002) we find it can satisfactorily be approximated by a normal distribution (fig. 2).

**Figure 2:**
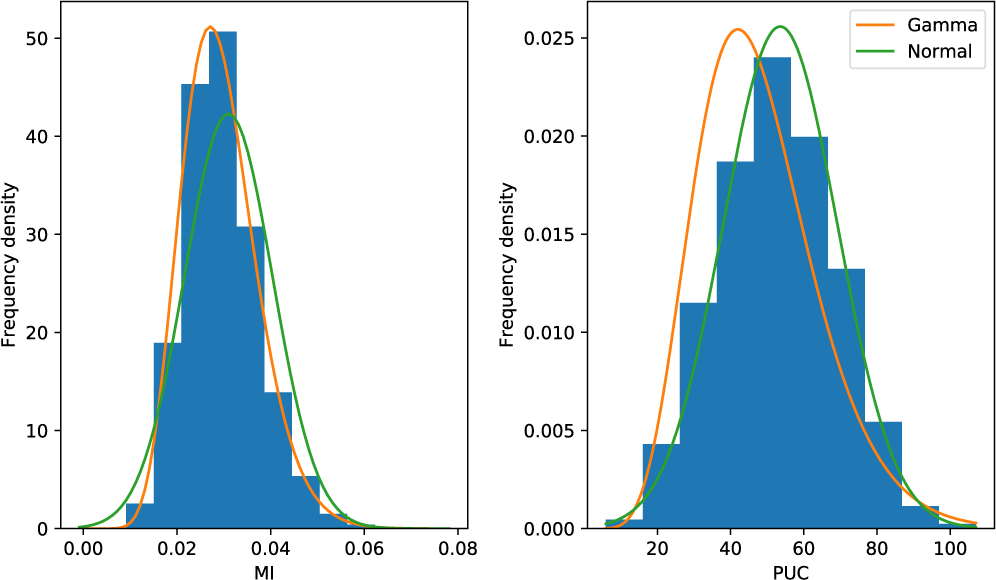
Distribution ofMI and PUC for all pairs from a randomly generated dataset with 100 genes and 1000 cells. The gene expression measurements are independent and normally distributed, and discretized using the Bayesian blocks algorithm. The lines show the null distributions fit by mode-matching, using all ofthe data, since it is all expected to be null-distributed.

We estimate empirical null distributions of MI and PUC using a regression-based mode-matching method outlined by (Efron, 2007) and extended for the exponential family by (Schwartzman, 2008). The method relies on the zero-assumption, that within a certain interval around the mode the contribution of *f_1_ (x)* is negligible; a density is fitted to a histogram of the MI (or PUC) scores falling within this interval. The zero-assumption interval and width of histogram bins must be chosen; we find that the method is fairly robust to both and more reliable than permutation and direct fitting methods (fig. S1).

**Figure S1:**
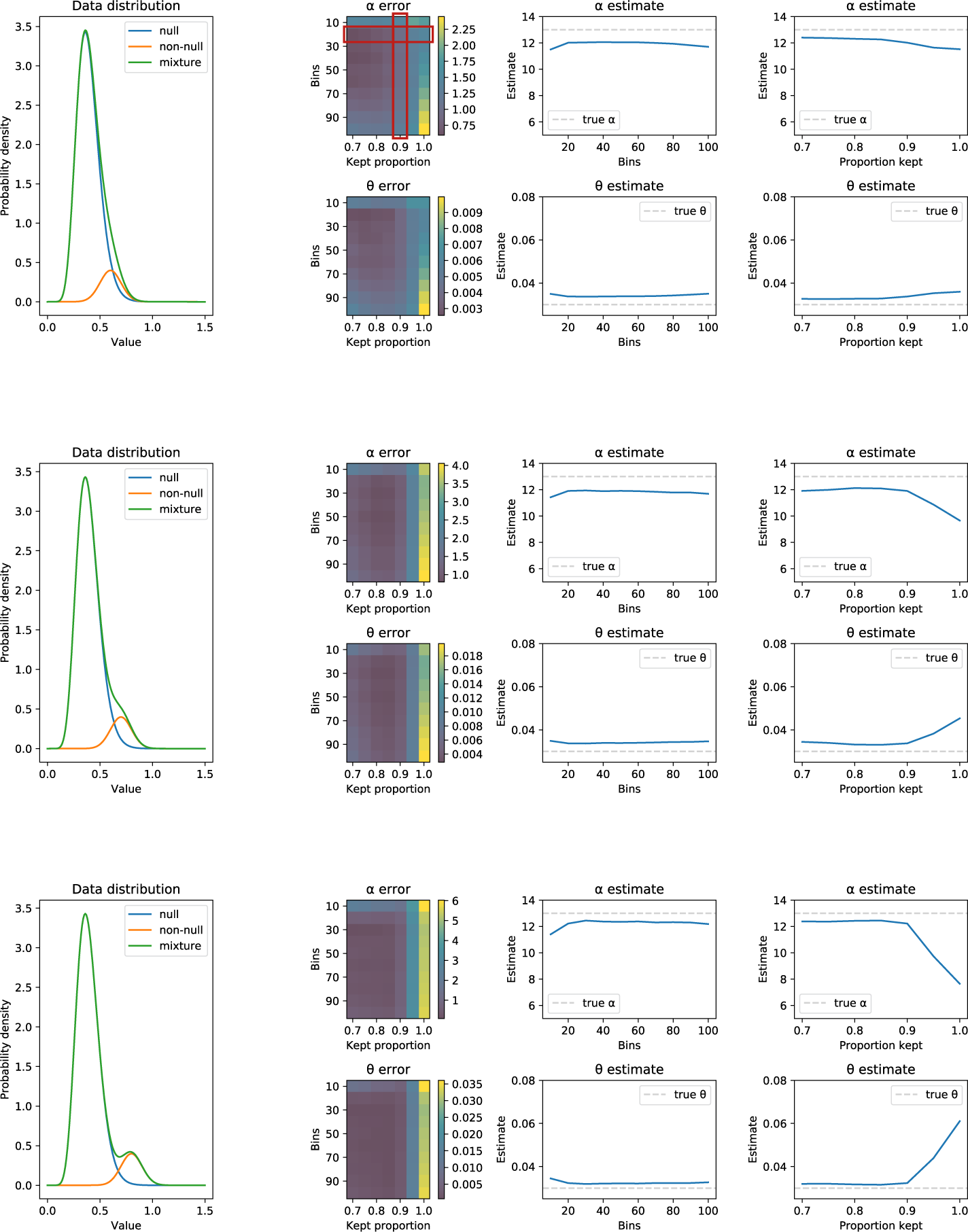
The effect of the number of bins and the zero-assumption interval on empirical null estimates for MI. We generated random test statistics using the mixture model in Eq. 3, assuming a gamma distribution for f_0_(x) and a normal distribution for f_1_(x), with π_0_ = 0.9 and *π_1_* = 0.1, with varying degrees of overlap between the distributions (line plots, left). We then used our mode-matching software to infer shape and scale parameters for f0(x) (true parameters were π_0_ = 13 and ɵ = 0.03 respectively). Errors are shown with (i) the number of bins and assumption interval varying together (heat-maps), (ii) the zero-assumption interval fixed at the lowest 90% of test statistics, and (iii) the number of bins fixed at 15 (line plots, right). Errors are on the whole small, except when the zero assumption is violated (i.e. the zero-assumption interval is too large).

We estimate the mixture density, *f(x),* by fitting a cubic smoothing spline to the full set of MI or PUC scores (Efron, 2004).

## Information Theoretical Network Inference using Prior Information

The mixture weights, or prior probabilities *π_0_* and *π_1_* = 1 - π_0_, have thus far been left unspecified. They represent our prior belief that any pair of genes in the network are unconnected or connected, respectively. We can (i) estimate these from the data (however, mode-matching methods tend to overestimate *π_0_* (Efron, 2004)); (ii) set *π_0_ =* 0.9 (a good approximation, e.g. for biological networks which tend to be sparse, and our default here, unless otherwise stated); or (iii) specify them from prior information.

Prior information, as described below (and elsewhere), comes in different forms, and ranges from specific information about pairwise interactions to information about global network characteristics (which in turn range from the expected number of edges in a network, to large scale hierarchical organization). *π_0_* and *π_1_* specify our prior belief about network sparsity, a global characteristic; however, with gene regulatory networks, we often have information about specific pairwise interactions (for example, we may know a particular transcription factor can bind to a particular target, in which case we may only require a weaker dependency in our transcriptomic data to infer an edge).

Unlike for conventional Bayesian inference, there is currently no straightforward mechanism within the empirical Bayes framework for incorporating such specific priors, so instead we introduce a *post hoc* adjustment to *π*_0_. For each potential edge *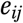* we may have different types of prior information, with different degrees of confidence, 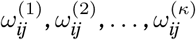 We can combine these, while simultaneously maintaining proper normalization, via

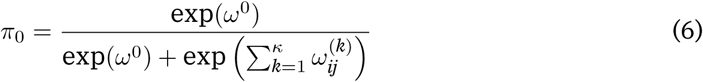

Where *ω_0_* does not depend on the particular edge; we find this functional form convenient and flexible, but this is not the only possible way of fusing different sources of prior information. With Eq. (6) we thus have for any candidate edge for which there is neither prior information for or against its presence,

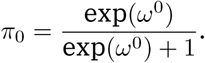

For most cases *π_0_* is determined by the value of *ω_0_,* which we set to 2.2 to give *π_0_ =* 0.9 (fig. S2).

**Figure S2:**
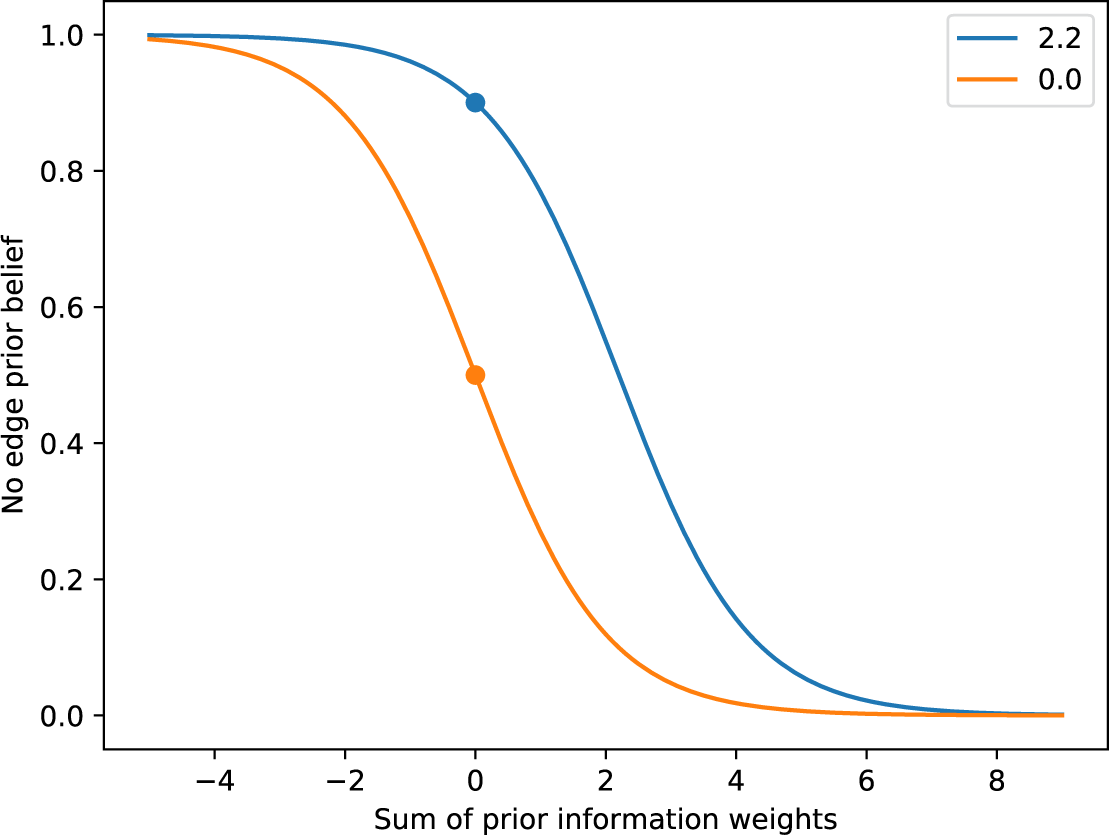
Prior networks using (A) the top 5% of edges from a network inferred using the PIDC algorithm from the population data (top 1% shown for easier comparison with single cell networks in fig. 5 and 6, right), and (B) protein-gene promoter binding interactions from the ESCAPE and CODEX databases. Assortativity of the three networks is 0.07 (population, top 5%), 0.06 (population, top 1%) and −0.07 (protein-gene promoter binding).

## *In silico* networks

### Data

We generated simulated datasets using a single-cell adaptation of the widely-used benchmarking tool GeneNetWeaver (Schaffter et al., 2011) as previously described (Chan et al., 2017). We simulated transcriptomic data for six networks with varying sparsity from *S. cerevisiae,* three with 50 genes and three with 100 genes (described in detail in (Chan et al., 2017)). Each dataset corresponds to 21 sample times and 2100 cells in total.

### Inferring Networks & Evaluation

The MI and PUC algorithms assign a score to all possible pairwise edges among a set of nodes, which are then transformed using Eq. 4 and 5. The output is a list of edges ranked from most likely to least likely to exist (as predicted by EB+IT).

We evaluate the accuracy of our ranked lists using the area under the precision-recall curve (AUPR), an appropriate measure for unbalanced binary classification problems such as network inference (Marbach et al., 2012). Higher AUPR scores correlate to better rankings, with more of the true edges ranked higher than false edges. To evaluate the performance for a particular network (instead of a ranked list), we calculate the precision, i.e. the proportion of predicted edges that are present in the true network.

### Significance testing

Creating a candidate network from a ranked list of information scores is problematic, because there is no formal framework for significance testing in an information theoretic context. A commonly-used thresholding method for information-based network inference is to include only the top *n%* of edges (Marbach et al., 2012; Chan et al., 2017). Since networks vary in connectivity, however, the most appropriate *n* is likely to differ between networks (e.g. in different organisms, or even cell-types); it is difficult to control network accuracy across studies using such a blunt approach. EB+IT seeks to alleviate this issue by assigning to each edge a*p*-value obtained from large-scale hypothesis testing. This exploits the information available in the data more fully, and allows us to obtain more consistent criteria by which to decide whether an edge is present or not. Among other things, this also provides some insight into the density or connectedness of a GRN (which is impossible if fixed fractions of edges are considered).

Figure 3 compares the spread in precision across different networks, varying the empirical Bayes threshold for including an edge in the network. Precision varies more predictably across networks and across algorithms when empirical Bayes is used, compared to keeping the top *n*%.

**Figure 3:**
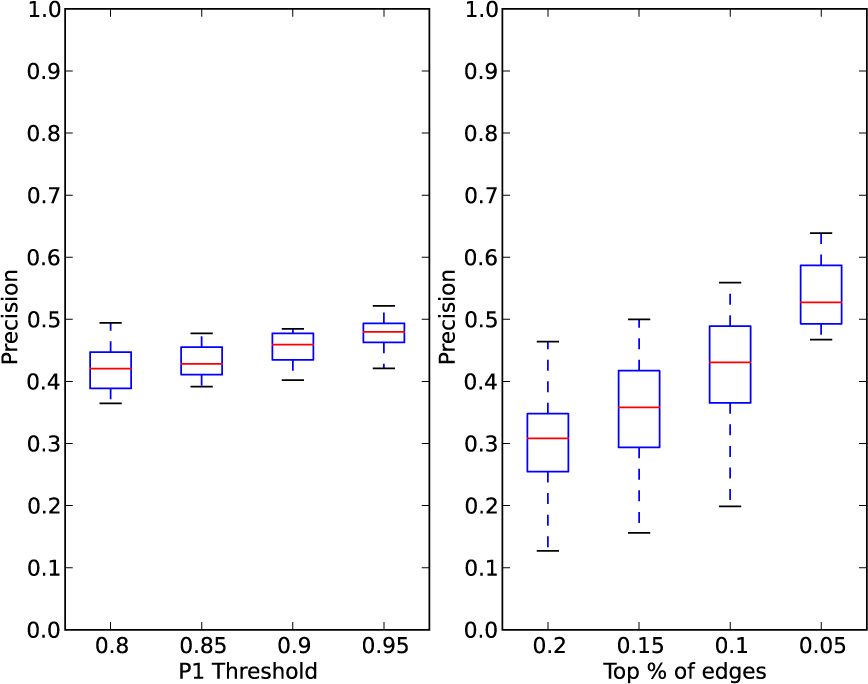
Variation in the precisión ofinferred networks across the six simulated datasets and the two inference algorithms, when using empirical Bayes thresholds (left) and the top n% ofedges (right). Precision is more consistent across different networks when the empirical Bayes score is used.

### Prior information

Since in the simulations the true networks are known, we can construct priors of known accuracy to test the effects of prior information. Our priors take the form of 1/0 values indicating the presence or absence of an edge. This set of values is generated using the methods mentioned in (Wang et al., 2013), taking some proportion, q, of random edges from the true network and then adding random incorrect edges to make a prior network of predetermined size. For example, a prior network with 100 edges and *q* = 0.3 consists of 30 true edges and 70 incorrect edges.

Figure 4 shows how AUPR varies with prior quality. Even relatively poor priors can improve the inference results, but very poor priors can worsen them. Without any prior information, PUC’s inferences are more accurate than those based on MI (and, correspondingly, prior information needs to be of a higher quality to improve PUC than MI networks).

**Figure 4:**
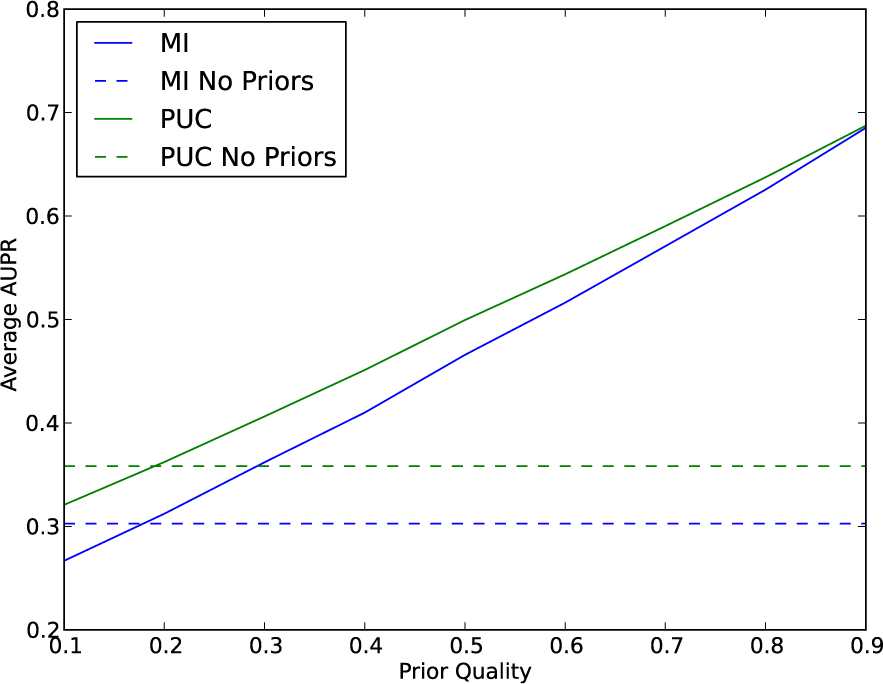
Mean AUPR across the simulated datasets whenpriors ofvarying knownprecision are used. Dashed horizontal lines indicate mean AUPR with no priors, for comparison.

## Biological pluripotency networks

### Data

We use single-cell transcription data from mouse pluripotent stem cells (PSCs) undergoing neural differentiation (Stumpf et al., 2017). PSCs were initially grown in fully defined conditions (2i) and differentiation was stimulated by growth in N2B27 media. 508 cells were quantified in total across 6 different time points for 168 hours.

### Evaluation

The difficulty with assessing the quality of a GRN inferred from experimental data is that we do not know the ground truth network; instead we rely on expert knowledge to assess plausibility of the inferred network. Current knowledge of GRNs is, of course, incomplete; nevertheless we have a set of observations and expectations that we can use to assess the inferred networks.

We know, for example, that biological networks are hierarchical and modular; we expect that a large fraction of interactions will be between nodes within the same module; we also know which genes should belong to the same module; and we have a good idea as to which modules are important in a given cell type or at a given developmental or physiological stage. An inferred network which does not capture these hallmarks of biological network organization must be viewed with suspicion. And we can use this indirect evidence together with the ability to detect known interactions to judge the quality of an inferred network.

We measure the ability of an inferred network to recover our knowledge of gene function using network assortativity (Newman, 2003). Briefly, if we associate each gene with a label (here we use the functional labels that were published alongside the original dataset, e.g. naive pluripotency, neuroectoderm), assortativity measures the extent to which nodes with the same label preferentially connect to each other, on a scale of −1 to 1, where 0 indicates no preference and 1 indicates that all edges connect similarly-labeled nodes. Assortativity also assigns a score to each label group, offering insight into which functional sub-networks are well-connected and also higher-level interactions between functional groups.

Since assortativity is influenced by network topology, (Bianconi et al., 2009) we also calculate the assortativity of 100 random graphs generated from each network using configuration models (Thorne and Stumpf, 2007) to determine whether the assortativity of the inferred network is higher than expected.

### Significance testing

In order to test the use of EB+IT for controlling the accuracy of networks inferred from experimental data, we inferred networks from the PSC data and calculated the assortativity, as the threshold for accepting an edge in the network was varied. Null distributions fit reasonably well for MI and PUC (fig. 5A). In agreement with the *insilico* networks, accuracy tends to increase as the threshold becomes more stringent with both MI and PUC, and accuracy is higher in general in the PUC networks (fig. 5B).

**Figure 5:**
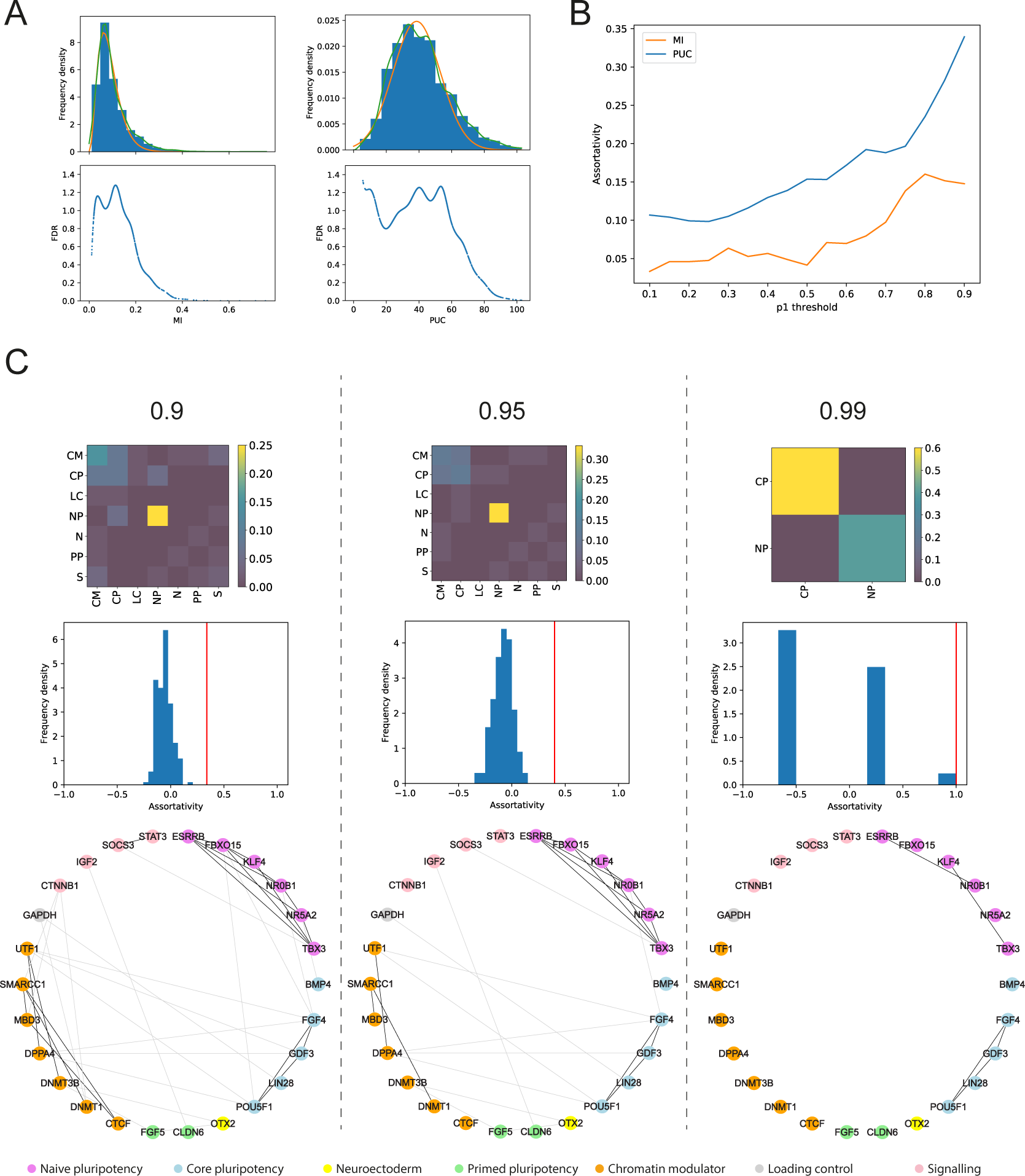
Application of EB+IT to PSC data. (A) Null and mixture distributions fitted to MI and PUC scores from the PSC data, and corresponding FDRs. A gamma distribution was fit to the MI scores and a normal distribution to the PUC scores, discretized into 20 bins and 15 bins respectively, using the lowest 90%for both scores. (B) Assortativity of PSC networks based on different empirical Bayes thresholds, using MI (orange) and PUC (blue). (C) Features of networks obtained from PSCs using the PUC measure, at empirical Bayes thresholds of 0.9, 0.95 and 0.99. Heat-maps show the assortativity ofeach network bygroup. Histograms show the distribution of assortativity when each network is randomly reconfigured 100 times using random configuration models (red lines indicate the assortativity of the inferred network). Network plots show connectivity between functional groups, with node color indicating function label and darker edges between similarly-labeled nodes.

We consider three PUC networks with thresholds of 0.9, 0.95 and 0.99 (for MI networks see fig. 6). The network with the highest threshold has an assortativity of 1.0, involving core pluripotency and naive pluripotency genes. We detect experimentally validated direct interactions, including Pou5f1-Fgf4 and Esrrb-Nr0b1, as well as new interactions that are strong candidates for future experimentation. However, with a 0.99 threshold there are very few edges. Reducing the threshold to 0.95 yields a larger network that captures the extensive interaction among naive pluripotency components in addition to genes involved in chromatin modulation (fig. 5B and C). Assortativity is lower, reflecting expected interactions between primed pluripotency and signalling, and, presumably, the lower FDR. These effects continue further at a threshold of 0.9 and now there is a strong chromatin modulation sub-network (fig. 5C).

**Figure 6:**
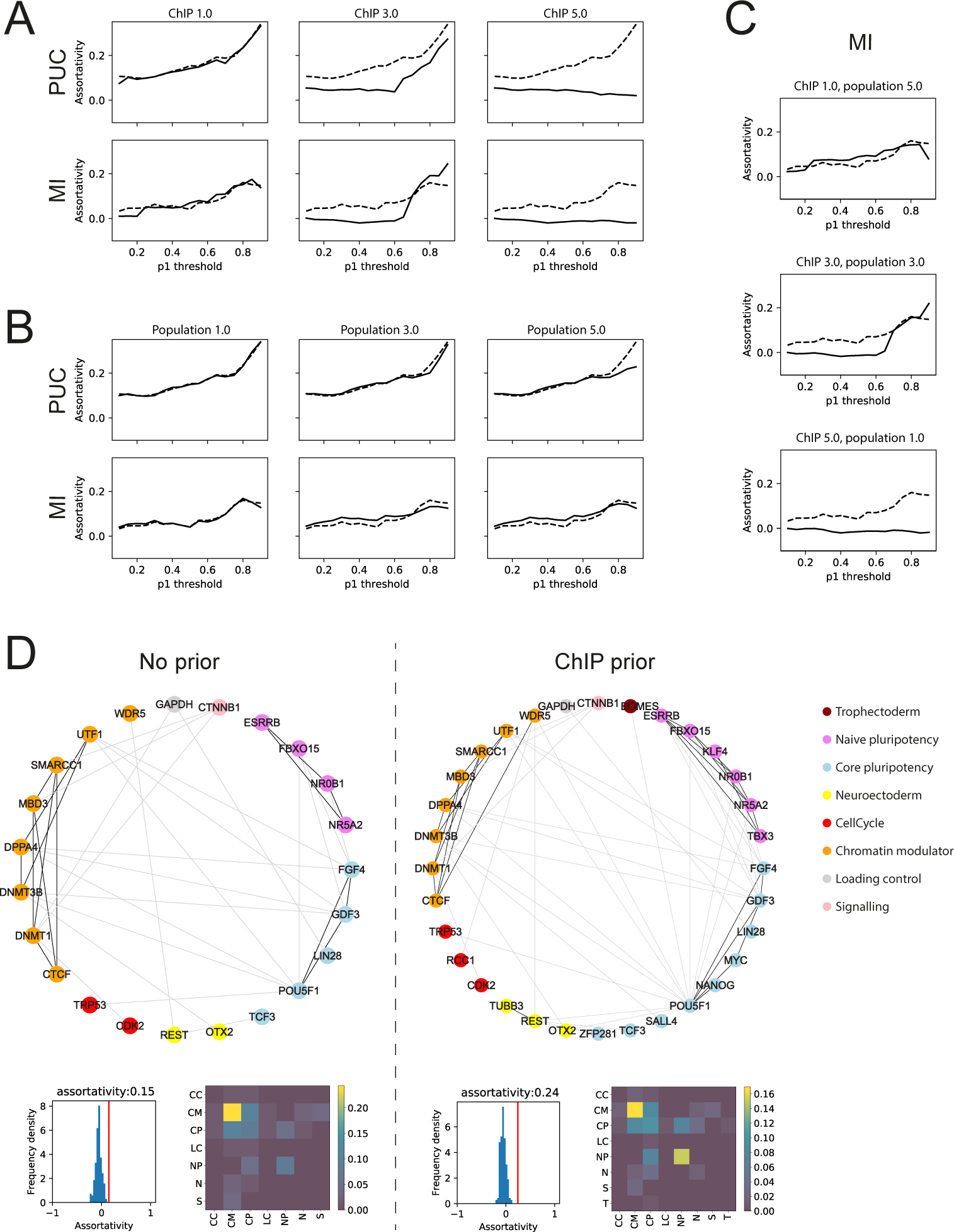
The effect of prior Information on networks inferredfrom PSC data. Assortativity as threshold is varied using (A) ChIP, (B)population and (C) both types ofpriors, with different weights. Dotted lines show assortativity when no priors are used. (D) Networks inferred using MI at an empirical Bayes threshold of0.9, with no priors and with ChIP priors with weight 3.0, and corresponding randomnetwork histograms and assortativity heat-maps.

### Prior information

We considered two types of prior information: population transcriptomic data produced alongside the single cell data (Stumpf et al., 2017), and protein-gene promoter binding interactions from the ESCAPE and CODEX databases, collated from chromatin immunoprecipitation (ChIP-chip/seq) studies (Xu et al., 2013; Sánchez-Castillo et al., 2014). The ChIP priors provide information for and against pairwise interactions, scored 1 and −1 respectively. Data are limited to the proteins tested thus far in PSCs and as such most interactions were untested; these were given a score of 0. Population networks were ranked lists, as discussed above, so we scored the top 5% of interactions 1, the lowest 5% −1, and the rest 0. Different prior weights were tested (see also fig. S2).

None of the priors appear to improve networks inferred using PUC (fig. 6A and B), and the ChIP priors with higher weights in fact worsen performance according to the criteria set out above. There is more room for improvement with the MI networks, however; for example, assortativity increases over the highest thresholds when low-medium weight ChIP priors are used (fig. 6A), though at the cost of lower assortativity at lower thresholds. A marginal improvement over most thresholds can be gained using high-weight population transcriptomic priors (fig. 6B), and by combining these with low-weight ChIP priors (fig. 6C).

We do not know the precision of prior information gained from experimental data; however, these results suggest that, according to our metrics, single cell data are more informative of functional relationships at a cellular level between genes than population transcriptomic or ChIP data (see also fig. S3). The slight improvement seen in the MI networks when prior information is used (albeit when carefully weighted) suggests that these types of experiments offer some useful information about the behavior of functional relationships that cannot be discovered by MI in single cell data. MI finds the chromatin modulation network but fails to discover much of the naive pluripotency network until the ChIP priors - which are primarily sourced from ChIP experiments of pluripotency transcription factors - are used (fig. 6D). On the other hand, the PUC measure appears to be sufficiently powerful to recover this information from single cell data alone: it discovers the naive pluripotency and chromatin modulation networks without prior information, and using prior information either has no impact or is detrimental to network quality.

**Figure S3:**
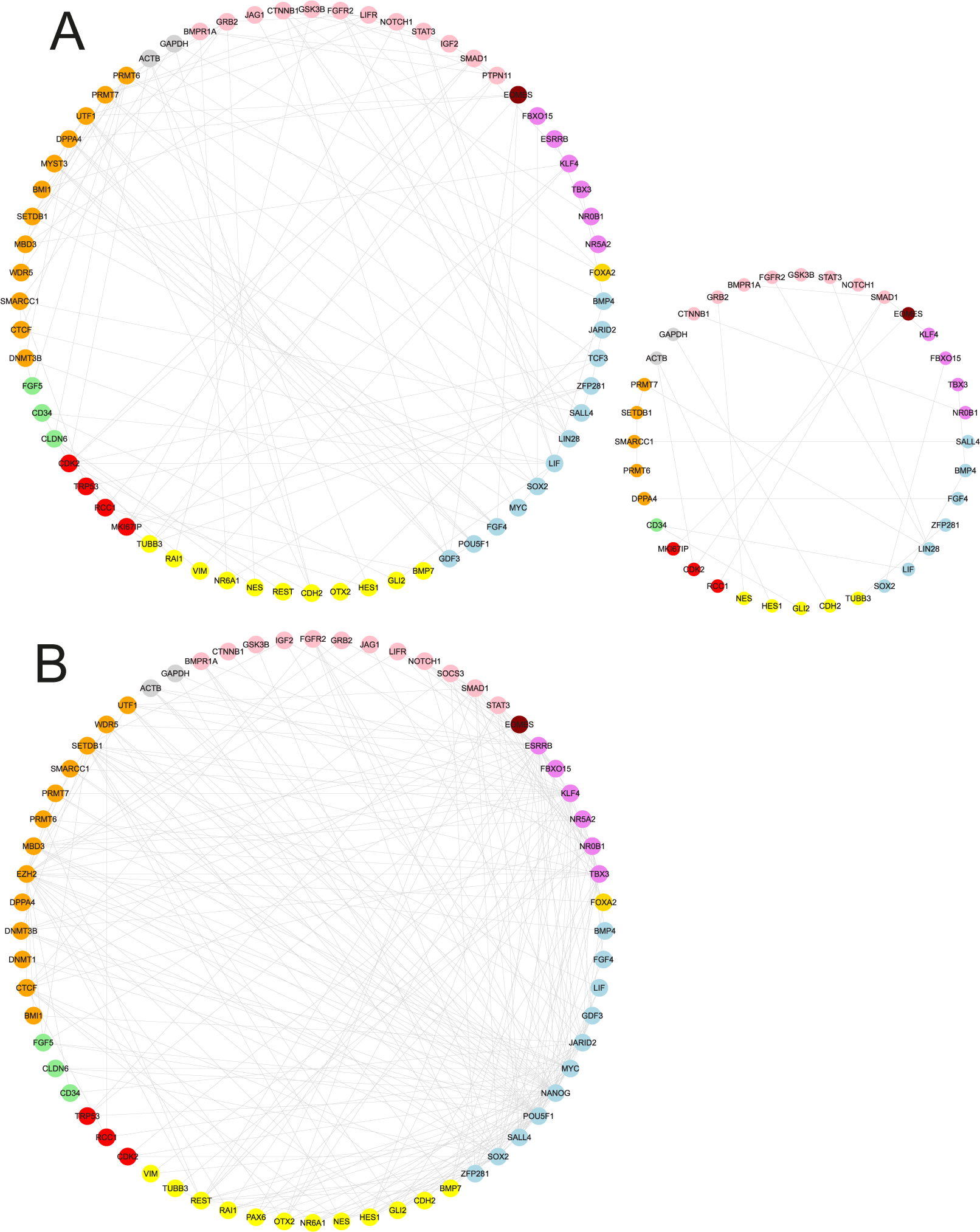
Prior networks using (A) the top 5% of edges from a network inferred using the PIDC algorithm from the population data (top 1% shown for easier comparison with single cell networks in fig. 5 and 6, right), and (B) protein-gene promoter binding interactions from the ESCAPE and CODEX databases. Assortativity of the three networks is 0.07 (population, top 5%), 0.06 (population, top 1%) and −0.07 (protein-gene promoter binding).

## Discussion

Here, we have introduced EB+IT, a method for applying formal large-scale hypothesis testing to information theoretic network inference, which also allows prior information about specific edges to be incorporated. Using *in silico* test cases, we show that EB+IT provides greater consistency in accuracy between networks inferred from different datasets, compared to the traditional approach of taking an arbitrary percentage of the highest scoring edges (fig. 3), and we offer guidance for when prior information might be helpful (fig. 4 and 6). We demonstrate these principles in a biological setting by inferring and analyzing networks from single cell transcriptomic data from differentiating mouse pluripotent stem cells; using EB+IT to control network accuracy, we propose a small number of candidate gene regulatory relationships, and discover higher-level interactions between sub-networks (fig. 5).

Several recent studies have inferred networks from single cell transcriptomic data, often as part of a wider analysis. These networks are inferred for a variety of purposes, including some that require a high edge confidence, such as identifying direct relationships for experimental validation (Moignard et al., 2013) and inferring detailed mechanistic models for making simulation-based predictions (Moignard et al., 2015; Matsumoto et al., 2017). Others examine the higher-level structural properties of the network and can therefore tolerate individual edge inaccuracies, such as supporting evidence of different cell states identified through clustering (using either global network connectivity (Pina et al., 2015) or the connectivity between groups of genes with known functions (Moignard et al., 2013; Stumpf et al., 2017)) and hypothesizing about particular genes’ roles based on their node characteristics within the network structure (Stumpf et al., 2017). With no mechanism for controlling network accuracy, it is difficult to use information-theory derived networks for purposes that require a high confidence (without making assumptions about the expected number of significant edges), and impossible to examine properties such as changes in global network activity (as in (Pina et al., 2015)), since the number of regulatory interactions is defined arbitrarily *apriori.*

Although single-cell data pose considerable challenges in the form of both technical noise and biological confounding factors (e.g. transcription stochasticity, population heterogeneity, cell cycle influences, etc.), they potentially offer a number of advantages over population-level datasets; these include larger sample sizes, and insights from (and into the potential causes of) cell-to-cell variability (Munsky et al., 2009). Here we find that single cell data are on balance more informative about the interactivity of functionally related gene groups than population-level transcriptomic or ChIP-chip/seq data (fig. 5 and S3). Using EB+IT we improve MI networks using ChIP-chip/seq priors, but find that PUC infers even more accurate networks from single cell data alone, suggesting that there are useful statistical signals within these noisy datasets that can be detected with sufficiently powerful methods. While this is encouraging for future single cell studies, we are unlikely to gain a full understanding of gene regulatory networks using only transcriptomic data (invariant or rapidly fluctuating relationships are not expected to be captured in this way, for example), so an interesting prospect is the development of single cell ChIP-seq techniques (Rotem et al., 2015) and similar developments in single cell epigenomics (Clark et al., 2016); it is conceivable that such experiments could provide informative priors even for PUC-based network inference.

When inferring a network from a new single cell dataset for purposes that require accuracy to be controlled, we advise using EB+IT with the more powerful PUC algorithm. The most appropriate threshold will depend on the use-case, but we find that higher-level network structures become apparent when using a threshold of 0.9. In some cases it will be more appropriate to use MI; for example, when inferring a very large network (on the scale of several thousands of genes), PUC will become noticeably slower due to its cubic complexity. Although we would expect MI networks to be less accurate, we might be able to improve them using appropriate prior information. Here, we found ChIP-chip/seq priors offered more of an improvement than network priors based on population transcriptomic measurements, perhaps because of the complementary nature of ChIP and single cell data (whereas population and single cell transcriptomic data would be subjected to many of the same biases and limitations). We note that these priors, although they improve MI networks at low-medium weights, can become misleading or detrimental when weighted too highly. We further note that low-weighted priors tend to do no harm in general (fig. 6A and B); hence we advise using low prior weights, and if several separate sources agree on the same interaction, then the weight will increase accordingly due to Eq. 6.

There is a need for statistical methods that can fully exploit the larger, more complex datasets produced by ever-improving experimental technologies. PUC and the related PIDC algorithms analyzing single cell transcriptomic data are examples of this (Chan et al., 2017; Stumpf et al., 2017). EB+IT enables a more intelligent, controlled approach to biological network inference from multiple sources of data, bringing together recent work in multivariate information theory, empirical null estimation, and large-scale empirical hypothesis testing. Our open-source software package EmpiricalBayes.jl implementing this approach is available, along with instructions and test data, from https://github.com/ananth-pallaseni/EmpiricalBayes.jl.

## Availability of data and materials

The EmpiricalBayes.jl package is available from https://github.com/ananth-pallaseni/EmpiricalBayes.jl.

A Julia package for running the PUC and MI algorithms is available from https://github.com/Tchanders/NetworkInference.jl.

Network inference tutorials and simulated datasets are available from https://github.com/Tchanders/network_inference_tutorials.

## Acknowledgements

This work was supported by a Biotechnology and Biological Sciences Research Council (BBSRC) DTP Studentship to TEC, a BBSRC Future Leaders Fellowship to ACB and an Imperial College London Fellowship to KRM. We thank Gal Barel, Suhail Islam for computing support, and Ben MacArthur and Patrick Stumpf, as well as the members of the theoretical systems biology group for useful discussions.

